# Cardiac-Gated Spectroscopic Photoacoustic Imaging for Ablation-Induced Necrotic Lesion Visualization: In Vivo Demonstration in a Beating Heart

**DOI:** 10.1101/2022.05.23.492682

**Authors:** Shang Gao, Hiroshi Ashikaga, Masahito Suzuki, Tommaso Mansi, Young-Ho Kim, Florin-Cristian Ghesu, Jeeun Kang, Emad M. Boctor, Henry R. Halperin, Haichong K. Zhang

## Abstract

Radiofrequency (RF) ablation is a minimally invasive therapy for heart arrhythmia, including atrial fibrillation (A-fib), which creates lesions using an electric current to isolate the heart from abnormal electrical signals. However, conventional RF procedures do not involve intraoperative monitoring of the area and extent of ablation-induced necrosis, making the assessment of the procedure completeness challenging. Previous studies have suggested that spectroscopic photoacoustic (sPA) imaging is capable of differentiating ablated tissue from its non-ablated counterpart based on PA spectrum variation. Here, we aim to demonstrate the applicability of sPA imaging in an in vivo environment, where the cardiac motion presents, and introduce a framework for mapping the necrotic lesion using cardiac-gated sPA imaging. We computed the degree of necrosis, or necrotic extent (NE), by dividing the quantified ablated tissue contrast by the total contrast from both ablated and non-ablated tissues, visualizing it as continuous colormap to highlight the necrotic area and extent. To compensate for tissue motion during the cardiac cycle, we applied the cardiac-gating on sPA data, based on the image similarity. The in vivo validation of the concept was conducted in a swine model. As a result, the ablation-induced necrotic lesion at the surface of the beating heart was successfully depicted throughout the cardiac cycle through cardiac-gated sPA (CG-sPA) imaging. The results suggest that the introduced CG-sPA imaging system has great potential to be incorporated into clinical workflow to guide ablation procedures intraoperatively.

## 1. Introduction

Atrial fibrillation (A-fib) is the most common type of heart arrhythmia [1]. In A-fib, the left and right atria receive chaotic bioelectronic signals, causing an irregular and often rapid heart rate. A-fib can induce blood clots in the heart and increase the risk of stroke, heart failure, and other heart-related complications. Between 2.7 and 6.1 million patients are currently affected by A-fib in the United States. According to the Centers for Disease Control and Prevention (CDC), this number is estimated to rise to 12.1 million by 2030 [2]. The current standard of care for drug-resistant A-fib is to use percutaneous catheter ablation therapy, which includes radiofrequency (RF) ablation. It aims to create lines of targeted tissue necrosis surrounding the pulmonary veins by creating lesions that protect the heart from abnormal electrical signals originating from the pulmonary veins that trigger A-fib. However, the lack of intraoperative evaluation of ablated tissues can lead to inadequate or excessive ablation. A-fib recurrence within the first year after the operation is common and is mostly associated with inadequate ablation [3]. Excessive ablation could lead to steam pop, a relatively infrequent (0.1% to 1.5%) but potentially severe complication [4,5]. Steam pop has been associated with embolic strokes, cardiac perforations, and ventricular septal defects. While imaging modalities such as magnetic resonance imaging (MRI) and computed tomography (CT) are clinically used to assess ablation lesions, neither modality provides real-time feedback, and their intraoperative usage is still in its infancy. Monitoring the necrotic lesion formation intraoperatively is critical to prevent excessive or incomplete ablation.

Photoacoustic (PA) imaging is an emerging imaging technology based on laser-generated ultrasound (US), which depicts the tissue’s optical absorption [6–8]. Different materials possess unique spectroscopic characteristics, and these can be used to characterize different types of chromophores [9,10]. The imaging modality has been widely used for vascular mapping [11– 15], tumor detection [16–18], and oxygenation mapping [19,20]. The implementation of PA imaging has been investigated in intravascular imaging for vulnerable plaques detection [21– 25], as well as for intraplaque hemorrhages in carotid artery plaque [26,27]. PA imaging has also been used for cardiac arrhythmia. Graham *et al*. reported a PA-based localization of the cardiovascular catheter, where they attached an optical fiber to generate a beacon PA signal from the tip of the catheter [28]. The same group also reported a pipeline implementing PA-guided cardiac ablation from navigation to ablation monitoring [29]. The monitoring of RF ablation using PA imaging was also reported in the previous works, where it was noted that the PA spectrum of ablated tissues differed from that of non-ablated tissues [30,31]. The ablated tissue contrast could be highlighted with respect to surrounding non-ablated counterparts through spectroscopic PA (sPA) imaging. Iskander-Rizk *et al*. demonstrated the real-time implementation of cardiac ablation monitoring by visualizing the intensity ratio of two-wavelength PA data in an *ex vivo* setting [32–34]. Although the abovementioned works made a substantial contribution to the feasibility of sPA imaging for ablation lesion evaluation, the assessment of lesion continuity and ablation extent can be challenging. It is difficult to determine whether weak tissue contrast is caused by insufficient light illumination or the presence of viable tissue, as both are present in the same image. Effective visualization of the lesion boundary between ablated and non-ablated tissues is vital for providing accurate feedback, so it can guide the intraoperative ablation procedure. More importantly, the cardiac applicability of the sPA imaging in an *in vivo* environment has not been demonstrated. The dynamic heartbeat motion can negatively impact sPA imaging performance. Additionally, the interference from other contrast sources, such as blood, is not negligible in certain configurations of implementing PA-guided cardiac ablation. While a fiber will be in full contact with tissue when the fiber is integrated with the ablation catheter [35], accessing a wider PA imaging field or using non-contact ablation introducers (i.e., laser [36,37]) requires illumination with a distance from the tissue, compromising the tissue contrast due to the blood absorption.

In this paper, we introduce a framework for mapping the extent of tissue necrosis, based on sPA imaging with cardiac gating incorporated, to explore the applicability of the sPA ablation monitoring in an *in vivo* environment. We computed the degree of necrosis, or necrotic extent (NE), by dividing the quantified ablated tissue intensity by the total intensity from both ablated and non-ablated tissues, visualizing it as a continuous colormap to highlight the necrotic area and extent. To compensate for tissue motion during the cardiac cycle, we applied the cardiac-gating on sPA data, based on the image similarity. The concept is validated in the *in vivo* swine model, where it allowed for the assessment of the ablation-induced necrotic lesion on the beating heart surface.

This paper first introduces the methodology for computing NE mapping and an image similarity-based cardiac-gating algorithm for cardiac motion compensation. Then, the experimental setup and animal preparation for *in vivo* validation are described. The results present the imaging performance compared with the control groups and the system tolerance with limited PA data to evaluate the potential for real-time implementation. Finally, the findings from the study and limitations are discussed.

## 2. Materials and Methods

### 2.1 Acronyms Necrotic Extent Mapping

This paper introduces a necrotic region mapping method, quantifying the extent of ablation-induced necrosis with respect to the non-necrotic tissue. After decomposing the intensity distribution of two myocardium states, NE will be quantified based on the contribution from the necrotic tissue intensity with respect to the total intensity combined with that of non-necrotic tissue.

Biomedical materials each have unique optical absorption characteristics consisting of PA spectra. The measured PA signals consist of a combination of signal intensity from multiple chromophores and contrast agents. Decomposing each material contribution and forming a sPA image can be conducted through spectroscopic decomposition (also known as material decomposition or spectral unmixing) [38]. Such a technology has been well explored in blood-oxygenation mapping [39], neurovascular mapping [40] and contrast agent enhanced imaging [38]. Based on the assumption that the received PA spectrum consists of a linear combination of multiple chromophores, the contribution from each contrast source is estimated as Eq. (1) [38].

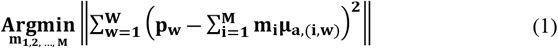

Here, *p* is the measured PA spectrum, *i* is the contrast source number, *M* is the number of absorbers, *μ*_*a*_ is the absorption spectrum of contrast source *i* at the wavelength *w*, and *W* presents the number of wavelengths used. *m* is the estimated composition of the contrast source.

In this proposed work, we first implemented spectroscopic decomposition to distinguish the contribution of the intensity distribution of the ablated tissue from that of non-ablated tissue. With the reference PA spectra from ablated and non-ablated tissues as the input, the intensity distribution from two tissue types is decomposed. Fig.1 shows the PA spectra used in this study, which were obtained from our previous work with the swine model [41]. An equivalent spectral trend is observed in the work reported by Dana *et al*. [31].

**Fig. 1.**
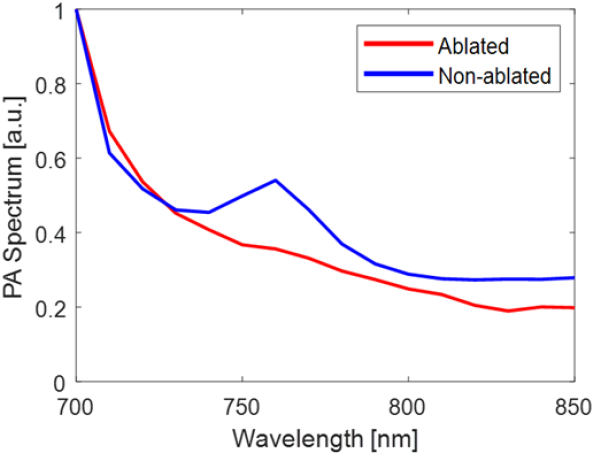
The photoacoustic (PA) spectra from ablated and non-ablated tissues used for spectroscopic decomposition [41].

The second step of quantifying the extent of tissue necrosis is to estimate the contribution of the ablated tissue intensity distribution with respect to the combination of non-ablated tissue intensity based on the decomposed intensity distribution from both ablated and non-ablated tissue types. This is based on the assumption that non-necrotic tissue should not contain the intensity shown for ablated tissue, and fully ablated necrotic tissue should not present this for non-ablated tissue. The NE metric presents the intensity ratio between ablated tissue intensity (*m*_*Ab*_) and the summation of ablated tissue intensity with non-ablated tissue (*m*_*N-Ab*_), as shown in Eq. (2). Such a mapping method is capable of capturing the transition between two tissue statuses.

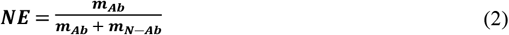

In addition, it is necessary to account for the potential occurrence of insufficient light illumination during scanning and the presence of background noise. In such cases, the measured PA spectrum will contain signals non-specific to neither ablated nor non-ablated tissue spectra, which could affect the estimation accuracy of both ablated and non-ablated tissue intensities. Therefore, we regard those non-specific signal spectra as residual and filter out the pixels that contain high residual contrast with respect to the total PA intensity, so as to only visualize the contrast source with higher confidence of NE mapping. The symbols used correspond exactly to those defined in Eq. (1).

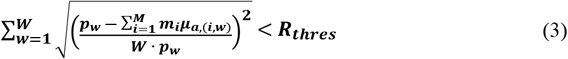

Here, *R*_*thres*_ is the residual threshold value which is set at 0.2 in this work based on the measured noise ground with the imaging device. This threshold enhances the tissue contrast in the scanned image by differentiating the low-intensity PA signal generated by the tissue from noise signals. The output NE map is displayed with a continuous colormap to show the transition of tissue necrosis in an intuitive visual sense, where red indicates ablated, and blue indicates non-ablated tissues, respectively. The NE value of 0.2 was set for defining the boundary of the ablated region.

### 2.2 Cardiac-Gated Spectroscopic Photoacoustic (CG-sPA) Imaging

The imaging speed of sPA imaging is determined by factors such as the laser pulse repetition frequency (typically 10–20 Hz or at most 100 Hz) and the number of wavelengths used for spectroscopic decomposition. Additionally, the signal-to-noise ratio (SNR) of the PA signals in the *in vivo* environment is expected to be compromised due to the limited space for placing light delivery fibers and light absorption from blood, requiring frame averaging to enhance the SNR. Hence, a certain temporal duration of PA data is required for sPA imaging in an intracardiac environment. However, continuous drastic tissue motion, due to heartbeat, introduces severe motion artifacts and prevents direct data processing. This motivated us to introduce cardiac gating to enable *in vivo* sPA imaging of a beating heart. Specifically, we track and map the recorded PA data to a specific time point with respect to a cardiac cycle. The image similarity, with respect to the specific time point, is used to sort and align PA data, in which averaging filter and spectral decomposition, and NE mapping are applied.

To track the temporal variation and similarity of PA images over time, we computed the 2D cross-correlation coefficient of two PA image frames. The coefficient *R* was defined in Eq. (4) as a scalar value between 0 and 1, of which 1 suggests two input images are identical and 0 indicates the opposite. The variable *m, n* is the pixel number in the input 2D envelope detected PA images *A* and *B*, with 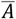 and 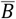 representing the mean value of *A* and *B* intensity.

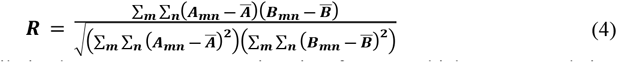

Based on the similarity between any two PA imaging frames, a high cross-correlation coefficient indicates the two cardiac PA images were taken when the cardiac tissue appears at a similar location in the field of view (FOV). Yet, a high coefficient *R* alone is not sufficient to extract a cardiac cycle because the tissue surface passes a similar location in the imaging FOV twice in a cardiac cycle, during contraction and relaxation. The contraction and relaxation data should be distinguished because the muscle status of the cardiac tissue could be different between them. This could have introduced additional noise in the spectrum when images were averaged. Therefore, we categorized the PA frames based not only on the image similarity of the current frame but also on the similarity of frames in a cardiac cycle.

To examine the similarity alignment of two frames with respect to a cardiac cycle, we computed the image correlation coefficient in the temporal axis. The temporal correlation coefficient was defined as *R*_*T*_ shown in Eq. (5), where *R* is the correlation coefficient defined in Eq. (4) that evaluated the similarity of PA image frame *f* with respect to a reference frame *f*^*r*^. The correlation coefficient of the frame number *i* in the current cardiac cycle was compared, as well as the frames (*i-j*) in the previous cardiac cycle, which includes *M* frames, where *j* is the frame number in the previous cardiac cycle. The *R*_*T*_ value reflects the image similarity and indicates the time point in a cardiac cycle. It allows the isolation of the signals from contracted and relaxed phases. The sorted high *R*_*T*_ frames guarantee both similarities in tissue position in the image FOV and alignment with respect to a cardiac cycle.

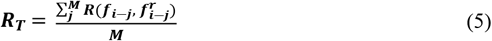

The aligned PA images, taken at the same timing in a cardiac cycle, were averaged to offer SNR-improved cardiac-gated (CG) PA images.

Fig.2 summarizes the workflow of signal processing for the recorded PA images, with the proposed CG-sPA imaging incorporating both cardiac gating and spectroscopic decomposition. The acquired PA channel data were first beamformed using a Delay-and-Sum (DAS) algorithm [42] and envelope detected, then sorted based on their PA excitation wavelengths. In each sorted wavelength frame set, a routine of the cardiac tissue surface motion was observed, and this was tracked with *R*_*T*_ to a defined reference frame in a particular cardiac cycle. At a specific time point in the cardiac cycle, the *R*_*T*_ of all the recorded PA frames was calculated with respect to the reference frame at this cardiac cycle. Frames with an *R*_*T*_ reading above the threshold (defined as 0.4 in our work) were selected and averaged. This thresholding definition allowed us to maximize the number of categorized frames while avoiding overlapping frames between each time point in the cardiac cycle. The same process was applied to all individual illuminated wavelength data. The SNR-enhanced outputs through averaging at each wavelength were combined as a CG-sPA data set for the following spectroscopic decomposition to distinguish ablated and non-ablated tissue spectrums. The NE map was computed as described in the previous section. The process was then repeated for the frames throughout a cardiac cycle.

**Fig. 2.**
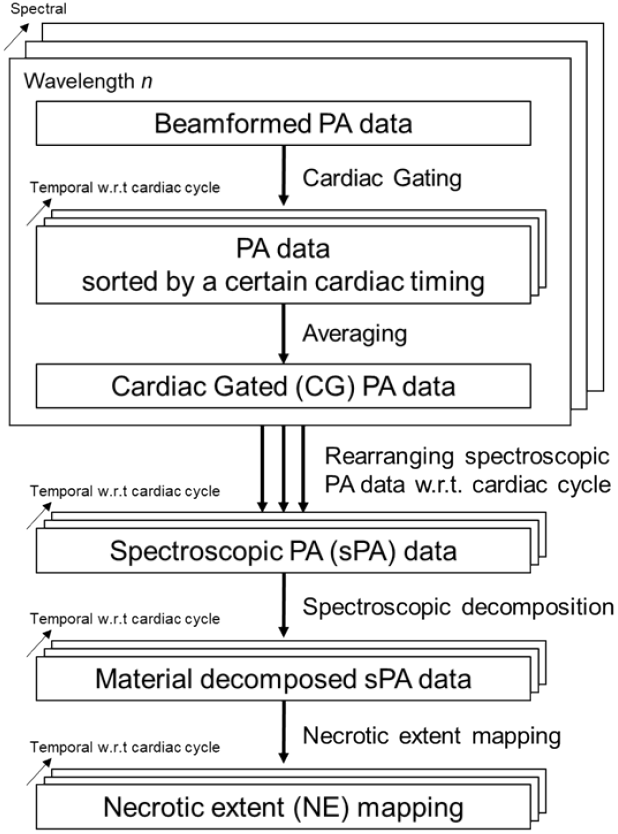
Cardiac-gated spectroscopic photoacoustic (CG-sPA) image-processing flowchart

## 3. Implementation

### 3.1 Experimental Setup

The PA imaging system setup used is shown in Fig. 3 (a). A 128-element linear-array US transducer probe (Philips ATL Lap L9-5, Philips, Netherlands) was used as the US receiver. The center frequency of the probe is 7 MHz with a bandwidth of 5 MHz to 9 MHz. The Verasonics Vantage system (Vantage 256, Verasonics, USA) was used as the data acquisition device. A Q-switched Nd:YAG laser (Q-smart 450, Quantel, USA) with an optical parametric oscillator (OPO) (MagicPRISM, OPOTEK, USA) was used, which is capable of generating a laser pulse at a wavelength range of 690–950 nm and at a repetition rate of 20 Hz with 5 ns pulse duration. A dual-head optical fiber bundle was used in this study to deliver light energy. Two fiber head were fixed on both side of the US transducer probe with tilted angle to focus light beam at approximately 10 to 15 mm depth.

**Fig. 3.**
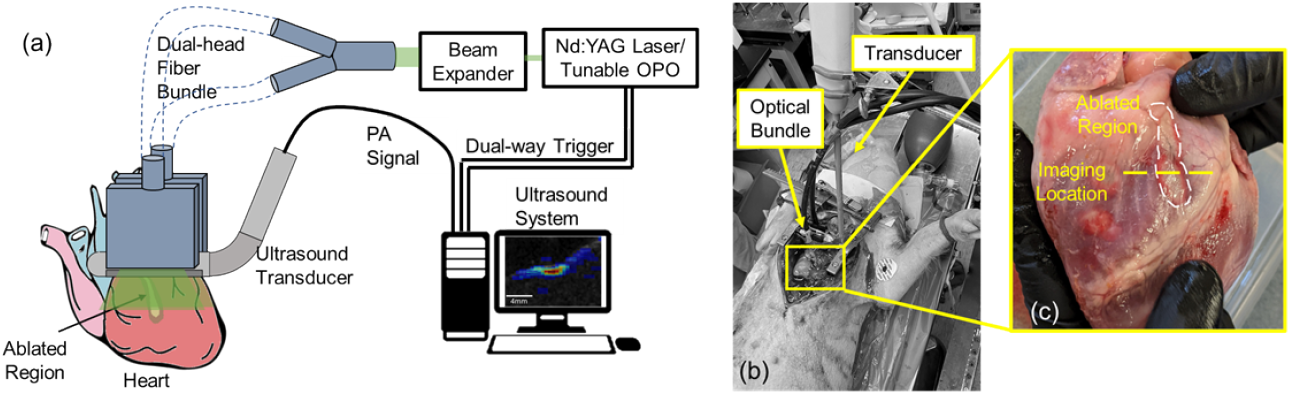
(a) The sketch of the imaging setup and system architecture. (b) *In vivo* cardiac photoacoustic (PA) imaging setup with swine model. (c) The post-procedure heart surface with the ablated region highlighted. The yellow dot-line indicates the imaging slice for necrotic detection.

We implemented two illumination modes for sPA imaging, as illustrated in Fig. 4. Three-wavelength repetitive illumination (3WL-Rep) allows a fast wavelength tuning between predefined sets of wavelengths. There is no tuning delay between wavelengths, and continuous scanning at a 20 Hz repetition rate is achievable. In this study, a 740 nm, 760 nm, and 780 nm wavelength laser were used and repetitively illuminated to acquire sPA images. This mode is suitable for achieving a high frame range of sPA imaging based on the limited number of wavelengths. The other mode, 16-wavelength individual illumination (16WL-Indiv), allows a range of laser wavelengths with a defined step size. A 700 nm to 850 nm wavelength laser was used with a 10 nm step. 64 frames of PA data were scanned at each wavelength. Having access to a higher number of wavelengths offers a more robust spectroscopic decomposition.

**Fig. 4.**
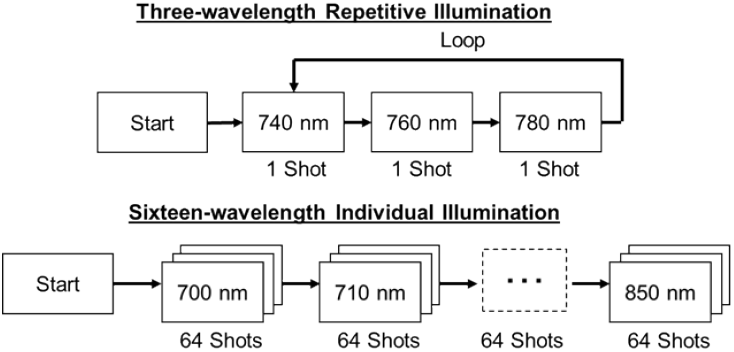
Two laser illumination modes: Three-wavelength repetitive illumination (3WL-Rep) is suitable to achieve a high frame sPA imaging, while 16-wavelength individual illumination (16WL-Indiv) offers a robust spectroscopic decomposition.

### 3.2 Animal Preparation and Study Procedure

A swine was prepared for a cardiac ablation procedure approved by the Johns Hopkins Animal Care and Use Committee (Protocol SW20M200, Approval Date: 7/12/2020). The swine was initially anesthetized with mechanical ventilation using a combination of ketamine and xylazine and maintained under sedation using 1–2% isoflurane. The swine was shaved, and its chest was opened to expose the heart for ablation and imaging. The subject was positioned supine, as shown in Fig. 3 (b), and was continuously monitored throughout the procedure. Ablation was performed using an irrigated 3.5 mm-tip ablation catheter (FlexAbility, Abbott, USA) on the part of the cardiac tissue surface, shown in Fig. 3 (c). The RF ablation was delivered at 30W with a temperature limit of 48°C. The irrigation speed was 17 mL/min. Under the *in vivo* environment in which the heart moves dynamically in a periodic manner, it is quite challenging to capture a particular ablated spot with a 2D ultrasound receiver because it easily goes off-plane. Therefore, we created a line-shaped ablation lesion and placed the ultrasound probe perpendicular to this line to minimize the out-of-plane uncertainty. The PA imaging probe was placed on the top of the cardiac tissue surface, ensuring the slice intersect the cross-sections of the line-shaped ablation lesion. (Fig. 3 (b)). Ultrasound gel (Aquasonic Clear Ultrasound Transmission Gel, Parker Laboratories, USA) was applied to couple the cardiac surface and transducer acoustically.

The experimental conditions are summarized in Table 1. Sham imaging procedures based on two illumination modes were performed at regions not containing the ablated tissue as the negative control. The imaging setup, illumination modes, and post-processing pipeline were identical to the ablation lesion imaging. Both imaging illumination modes were performed at the same ablated tissue plane. US scanning was performed at the same location before the PA imaging to acquire the anatomical information. The single-angle plane wave was used for US imaging. The US framerate was 20 Hz, which was the same as PA imaging to ensure the consistency. In the repetitive illumination (3WL-Rep) mode, wavelengths of 740 nm, 760 nm, and 780 nm were used, capturing the local spectrum peak in the non-ablated tissue at 760 nm. In the individual wavelength illumination (16WL-Indiv) mode, the wavelength range of 700 nm to 850 nm was used with a step of 10 nm.

**Table 1.** Experimental Conditions

|  | Experiment | Imaging Tissue Region | Ablation | US Scanning | Illumination Mode | Results |
| --- | --- | --- | --- | --- | --- | --- |
| #1 | Fast - Sham | Region 1 | No | Yes | 3WL-Rep | Figure 5, 6 |
| #2 | Robust - Sham | Region 2 | No | Yes | 16WL-Indiv | Figure 7 |
| #3 | Fast - Ablation | Region 3 | Yes | Yes | 3WL-Rep | Figure 5, 6 |
| #4 | Robust - Ablation | Region 3 | Yes | Yes | 16WL-Indiv | Figure 7 |

## 4. Results

### 4.1 In Vivo Visualization of a Beating Heart

The repetitive illumination of three wavelengths, 740 nm, 760 nm, and 780 nm, was performed for the fast scanning mode to distinguish two tissue types during the dynamic heartbeat motion. Fig. 5 shows the CG-sPA image using the repetitive illumination mode with a particular timing in the cardiac cycle. Approximately ninety seconds of sPA images are used for the CG algorithm. The image displayed the clear boundary of the cardiac tissue surface, aligning with the US image acquired at the same location and timing. From the spectral decomposed images, the ablated region was depicted on the expected ablated spot, while most of the PA signals in other regions were identified as non-ablated tissue. The NE map also showed a strong ablation-induced NE value in the ablated region. The ablated region containing ablated tissue signal intensity can be distinguished from the surrounding non-ablated tissue by the colormap in the NE map (Fig. 5). The average pulse energy was 46.1 mJ (min 25.38 mJ, max 82.82 mJ) in the repetitive illumination mode and 51.9 mJ (min 28.82 mJ, max 80.18 mJ) in the individual illumination mode across the selected wavelengths. The energy transmittance of the fiber bundle is approximately 40%. The estimated averaging surface fluence was approximately 10 mJ/cm2 based on the fiber bundle geometry and target distance. The approximate maximum fluence based on the recorded energy was 16.6 mJ/cm2, below the maximum permissible exposure (MPE) for safety. Considering the complex cardiac motion and environmental factors, the tissue surface fluence could be lower than the above estimation.

**Fig. 5.**
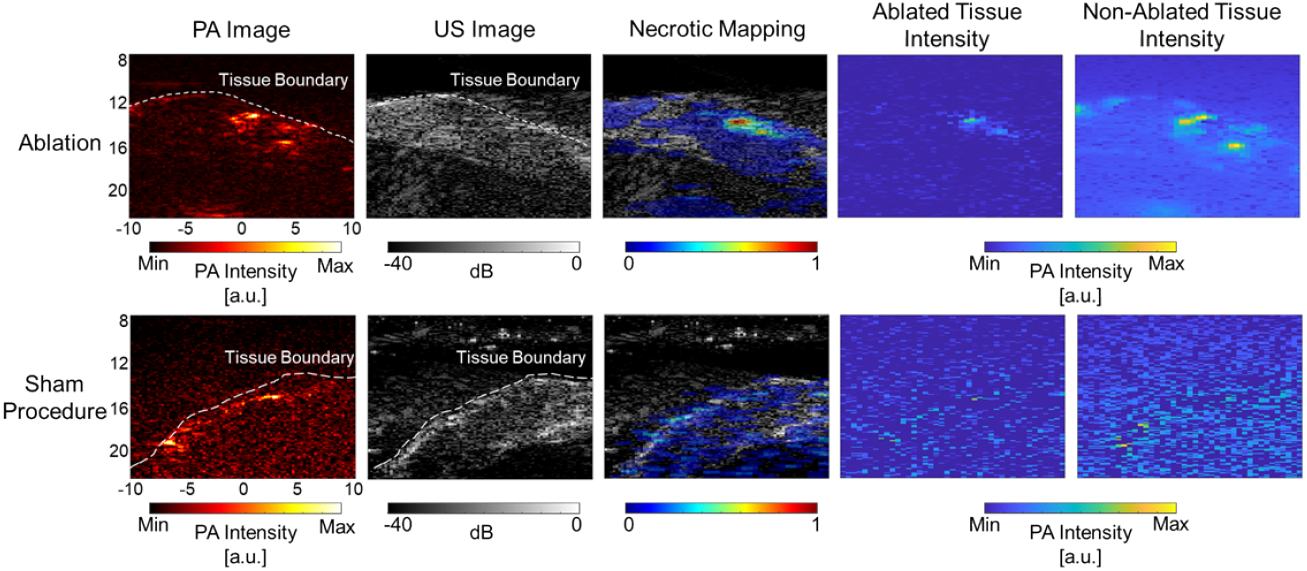
Cardiac-gated spectroscopic photoacoustic (CG-sPA) imaging results of the in vivo beating heart based on repetitive illumination. The ablated and non-ablated tissue intensity distribution maps show the outcome of the spectroscopic decomposition algorithm. The sham procedure was performed in a slice not containing ablated tissue. The displayed photoacoustic (PA) images were acquired with a 740 nm wavelength. [unit: mm]

A sham procedure was performed on a non-ablated region of the swine heart to validate the system specificity and confirm that the algorithm could identify the non-ablated tissue as non-ablated intensity. The same parameters used in the ablated tissue imaging session were used to acquire and process data. The results indicate that no obvious ablated signal intensity was detected. Most of the sPA signals were identified as the non-ablated tissue intensity. The NE mapping further confirms the non-ablated tissue identification.

To demonstrate the temporal variation of sPA-based necrotic mapping in the *in vivo* beating heart, Fig. 6 displays the CG-sPA images of an entire cardiac cycle, also available to view as Movie 1. The position of PA signal from tissue moved along consistently with the cardiac surface captured by US imaging, while NE mapping from CG-sPA imaging highlighted the ablation lesion at the center ablated area. The same tissue motion trends were observed in the sham procedure, with only the non-ablated tissue spectrum detected in NE mapping. Note that one heart cycle duration of 2.4 seconds is slower than a swine’s regular cardiac cycle duration. This is attributed to hyperthermia due to a time-lapse after the open-chest procedure.

**Fig. 6.**
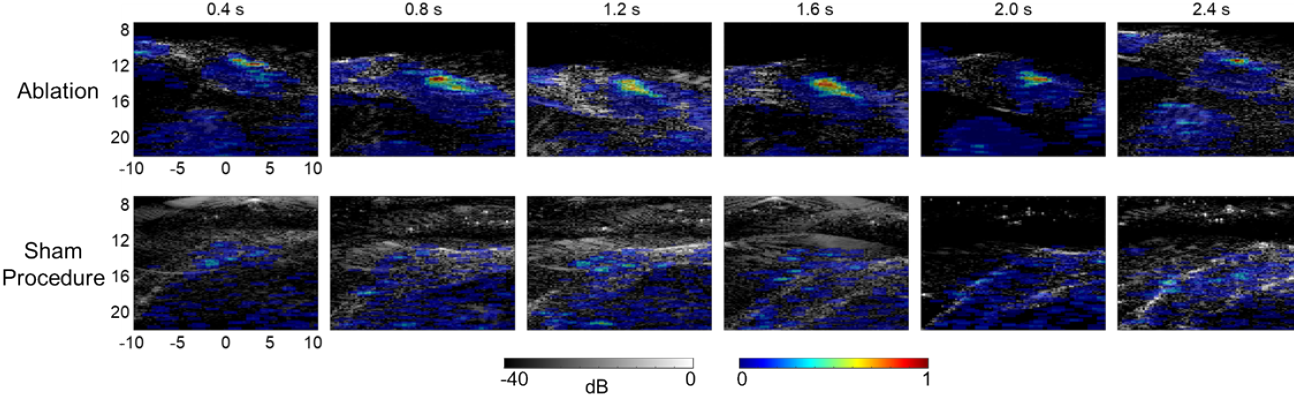
Necrotic extent (NE) mapping with repetitive illumination mode in one complete cardiac cycle, also available as Movie 1. [unit: mm]

### 4.2 Visualization with Individual Wavelength Illumination

We performed individual wavelength illumination scanning using 16 wavelengths, which captured a wide spectrum of the cardiac tissue. The wavelength range of 700 nm to 850 nm was used, matching the reported cardiac-ablation spectrum range [41]. A wider spectral range is expected to offer a more robust spectroscopic decomposition output by inputting more levels of wavelengths. To maximize the PA contrast from the recorded dataset, we calculated the number of frames available for averaging at each time point in the cardiac cycle and selected the time point containing the maximum number of available frames for presenting the NE mapping evaluation. The results from the individual wavelength illumination mode are displayed in Fig. 7.

**Fig. 7.**
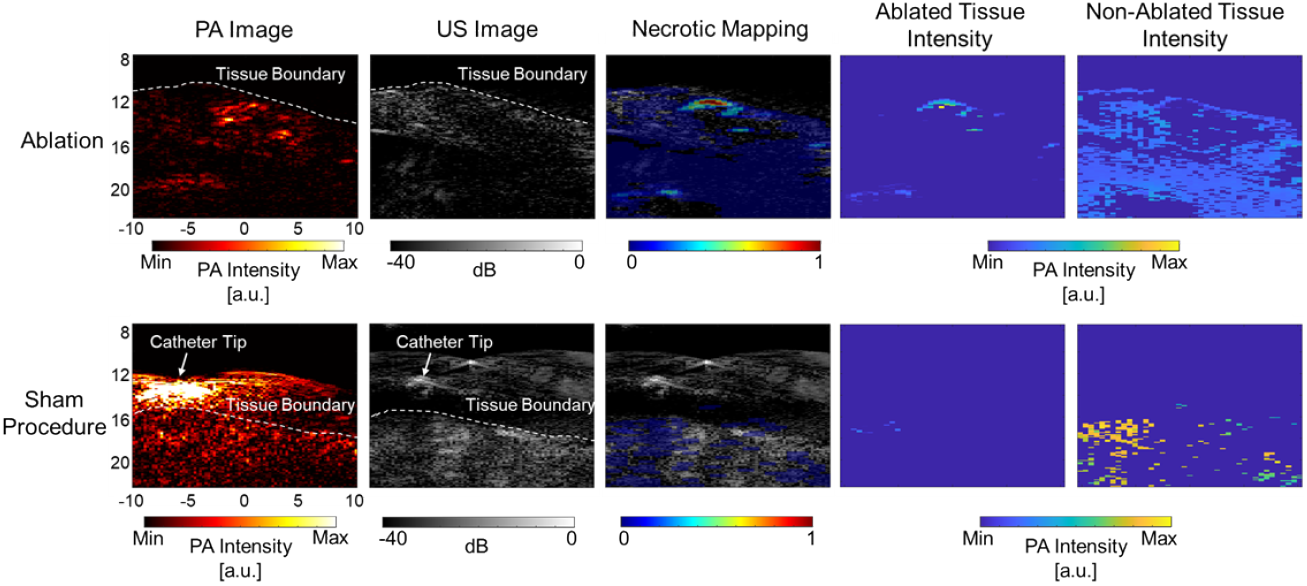
Cardiac-gated spectroscopic photoacoustic (CG-sPA) imaging results of the *in vivo* beating heart based on individual wavelength illumination. The ablated and non-ablated tissue intensity distribution maps show the outcome of the spectroscopic decomposition algorithm. The sham procedure was performed in a slice not containing ablated tissue. The displayed photoacoustic (PA) images were acquired with a 740 nm wavelength. [unit: mm] The displayed photoacoustic (PA) images were acquired with a 740 nm wavelength. [unit: mm]

This imaging session was performed immediately after the repetitive illumination imaging session so that the same location was imaged in equivalent conditions. The NE mapping based on individual wavelength illumination highlighted the same region depicted in the result from repetitive illumination. From the spectroscopic decomposed intensity distribution results, ablated tissue intensity was visible in the ablated tissue area, while non-ablated tissue intensity appeared throughout the tissue. The results from the sham procedure only capture non-ablated tissue intensity. Note that the sham procedures were performed at the different regions between repetitive illumination and individual wavelength illumination sessions.

### 4.3 Histological Validation

A histological comparison was performed between the necrosis region detected in CG-sPA imaging and the actual ablated tissue region of the heart to evaluate the accuracy of quantifying the geometry of the ablation-induced tissue necrosis. The cardiac tissue containing the ablation lesion was dissected after the swine was euthanized. The tissue sample was processed with 2,3,5-Triphenyltetrazolium chloride (TTC) to enhance the visibility of the ablation lesion. TTC is a marker of metabolic function and represents a reliable indicator of damaged areas in experimental models. It is a colorless water-soluble dye that is reduced by the mitochondrial enzyme succinate dehydrogenase of living cells into a water-insoluble, light-sensitive compound (formazan) that turns normal (non-ablated) tissue deep red. In contrast, damaged (ablated) tissue remains white, showing the absence of living cells, and thereby indicating the damaged region. As shown in Fig. 8, the measured dimension across the entire necrosis lesion was 5.51 ± 0.83 mm in width and 2.91 ± 0.43 mm in depth. In the NE mapping, the dimension of the ablation lesion scanned in the repetitive illumination scanning mode was 6.01 ± 2.35 mm in width and 1.79 ± 0.75 mm in depth. The dimension in the individual wavelength illumination scanning mode was 5.59 mm in width and 1.49 mm in depth, under an NE threshold of 0.2. Scanned lesion size is highly dependent on matching the measurement section with the imaged place, which was difficult to accomplish in this highly dynamic experiment.

**Fig. 8.**
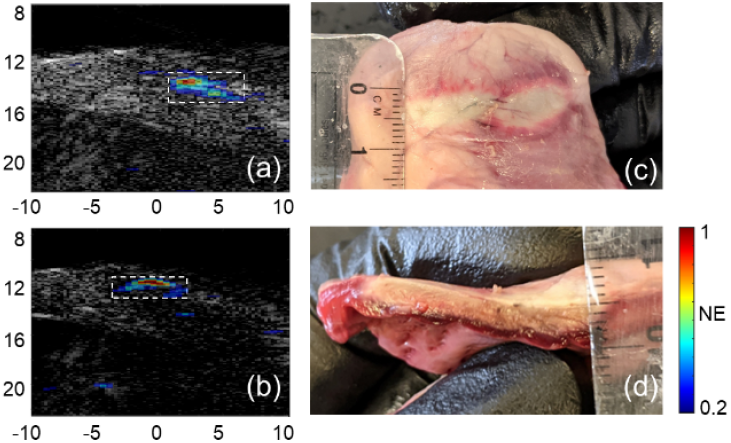
Histology analysis of the ablated tissue region. (a) Ablated region (white-dash rectangle) identified by the repetitive illumination scanning mode. (b) Ablated region (white-dash rectangle) identified by the individual wavelength illumination scanning mode. (c) Top view of the post-procedure processed ablated tissue for width measurement. (d) Side-view of the ablated tissue for depth measurement.

### 4.4 Evaluation of Imaging Duration

The proposed CG-sPA imaging algorithm uses the averaging filter to enhance the SNR and improve the outcome quality. However, using a long averaging filter compromises the system’s real-time imaging potential by requiring a large number of image frames as input. Therefore, we evaluated the impact of the averaging filter sample size on the system accuracy, in terms of sensitivity to ablated tissue identification (false-negative) and its specificity to non-ablated tissue identification (false-positive). This evaluation allows us to understand the required frame duration to grant acceptable performance for future applications. Fig.9 shows the NE mappings using data from varied durations of the PA recording time. The input data size is displayed in both the time duration of the sPA recording, as well as the mean number of frames for averaging after CG across wavelengths. With a greater number of sample frames used for averaging, the ablation lesion in the NE mapping got clearer as the noise reduced. The algorithm sensitivity was high with a smaller sample size where the necrotic region was detected. The small frame size presented low specificity due to high noise in the non-ablated region. The specificity was enhanced as the input data size increased.

Fig.10 presents the quantification of the NE map accuracy variation under different sample input sizes. The areas selected to quantify the NE value in the ablation region (ABL) and background non-ablated region (NAB) are shown in Fig. 10 (a). The PA SNR quantification of the tissue region is displayed in Fig. 10 (b). Due to the image layout, the noise region was set at the closest region above the tissue surface. The averaged NE value within these two marked regions was selected to represent and quantify the sensitivity and specificity of the necrotic lesion identification, respectively. Figures 10 (c-d) display the NE values with different input sample lengths (c) and with a different number of frames used in averaging (d), respectively. Both ABL and NAB region values started to be stabilized after using more than 20 frames for averaging, corresponding to more than 30 seconds of duration data. Further, the ratio of the mean NE values between the two regions was calculated to represent the ablated tissue detectability intensity distribution (Fig. 10 (e) and (f)). The NE ratio continuously increased with each increment of input data and reached saturation after around 25 to 30 frames of averaging.

## 5. Discussion

The proposed CG-sPA imaging in the repetitive illumination mode was able to visualize the ablation lesion in a beating heart. A boundary that was based on the NE value was able to isolate the ablation lesion from the non-ablated counterpart. The PA contrast was generally lower in the sham procedure compared to the condition that included the ablation lesion, which led to higher background noise. Despite the noise, the NE mapping did not suffer from a low SNR and presented non-ablated tissue distribution correctly. Additionally, the radiometric method used to calculate NE has the ability to account for variations in local fluence, ensuring that the NE mapping remains unaffected by the brightness of individual pixels in the sPA image. These observations demonstrate the robustness of the CG-sPA imaging and NE mapping. By processing NE mapping in the temporal axis throughout a cardiac cycle, it was observed that the ablation lesion moved with the heartbeat, corresponding to the shift of the cardiac tissue surface. While the lesion was consistently visible, the dimension of the ablation lesion varied between different timings in a cardiac cycle. This is attributed to two factors. First, the SNR of the ablation lesion may be temporally inhomogeneous because the light illumination dose varies as the position of the cardiac surface changes. Second, due to the nature of the heartbeat, the cardiac tissue remains still for a longer period in the relaxation phase. Therefore, a higher number of imaging frames for averaging was available at such a time. This resulted in a variation in the used number of frames for averaging in a cardiac cycle and affected the accuracy of the mapped NE value.

The results from the individual wavelength illumination scanning confirmed the presence of the ablation-induced necrotic lesion, aligning with the observation from the repetitive illumination scanning. The consistent visualization of the lesion supports the use of fewer wavelength levels in repetitive illumination compared to the thorough spectral illumination of 16 wavelengths in individual wavelength illumination. Regardless of the significantly higher number of wavelengths used, the decomposed intensity distribution between the two illumination modes were not considerably different because each wavelength only covers PA images for less than two cardiac cycles in the individual illumination mode, which results in less frame averaging in the CG-sPA processing. In fact, the tissue boundary was less clear in the sPA image in the sham dataset (Fig. 7). Still, it did not affect the non-ablated tissue identification in the NE mapping. Additionally, while the ablated tissue data for two imaging modes were collected at the same location, the cross-sectional B-mode images in Figs. 5 and 7 are not appeared to be similar because the images could capture slightly misaligned lesion slices due to the cardiac motion.

The histological analysis presented the necrotic lesion boundary identification capability with the CG-sPA imaging method. Both illumination modes showed a high agreement, especially in the lesion width measurement. The contrast degradation was more eminent in the depth axis due to the light scattering in tissue. Due to the complexity of the cardiac motion, it is difficult to set up the imaging probe perpendicular to the tissue surface and track the off-plane motion without robust mechanical motion compensation to move the transducer. With our presented imaging setup, it is hard to extract the one-to-one correspondence on the PA imaging location with the histology section for depth comparison. The imaging setup can only be localized on the line perpendicular to the lesion line to ensure it will be captured in the FOV. Furthermore, the ablation procedure of forming a line-shaped lesion was manually performed on a beating heart. Therefore, the consistency of the lesion width and ablation extent was hard to maintain across the ablated lesion. The potential localization error could cause a discrepancy in quantitative comparison between the PA measurement with histology.

The evaluation of the imaging duration for averaging offers guidance for optimal parameter selection in future real-time implementation. We quantified the accuracy of identifying ablated and non-ablated regions at various input data sizes, which indicated system tolerance to the short-duration data. The detection of the ablated tissue was achieved with less than 5 frames of input data, supporting the high sensitivity of the algorithm. In contrast, the non-ablated region identification was less straightforward due to the noise over a short duration of data (e.g. 5 frames). The false-positive detection was observed on both ablated and non-ablated regions with a short duration of data input. Overestimation of the NE value could lead to incomplete ablation during the procedure. The trend of suppressing a false-positive when input size increases match the observation in Fig. 9. Based on the imaging parameters, the false indication could be suppressed when more than 20 frames are averaged for each wavelength. A false-negative region is visible on the tissue surface in the 2.4 seconds time point of the sham result in Fig. 6 due to the smaller averaging window size (15 frames).

**Fig. 9.**
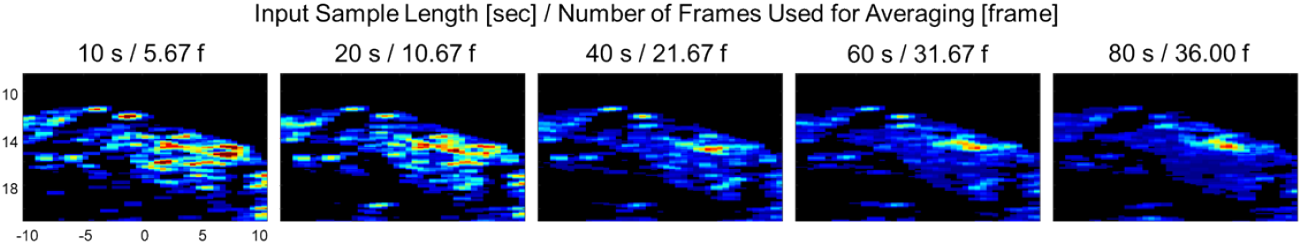
Necrotic extent (NE) mapping calculated with a varied input data length in terms of the time duration of the collected data and the mean number of frames used for averaging across each wavelength.

**Fig. 10.**
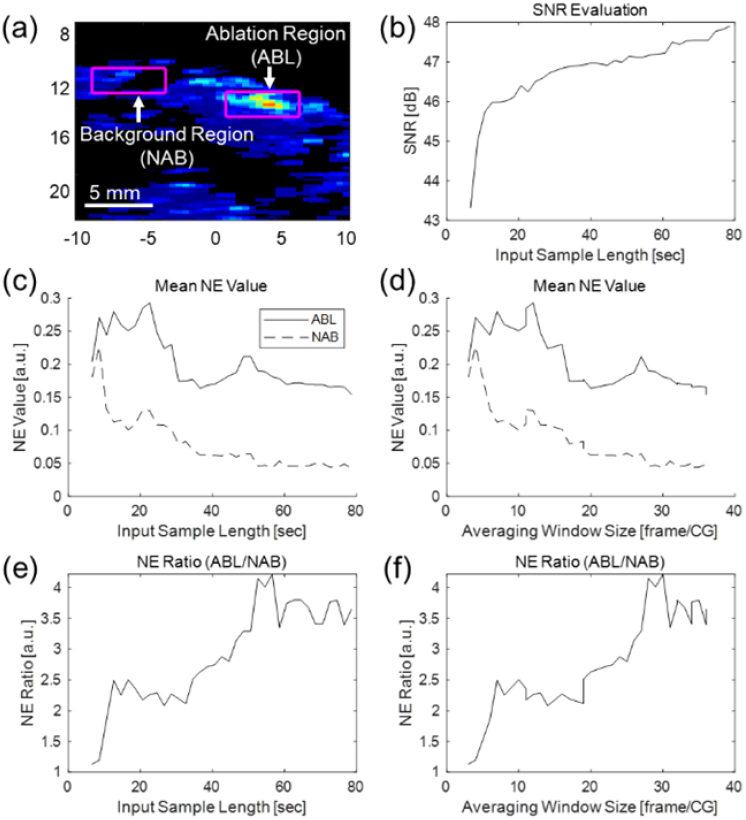
Quantitative evaluation of necrotic extent (NE) mapping based on the input data size. (a) A NE map based on 80 seconds of input data under repetitive illumination mode. Ablation (ABL) and background non-ablation (NAB) regions are marked. (b) The SNR improvement over varied input data duration. The mean NE value in the ABL and NAB regions changes over varied input data duration (c) and averaging window sizes (d). The ratio of NE values between two regions changes over varied input data duration (e) and averaging window sizes (f).

The proposed CG algorithm can serve as an add-on to the conventional ECG technique or an alternative when it is unavailable. Considering the PA implementation, achieving synchronization among multiple devices (like laser and ultrasound) is not trivial. The external triggering could contain delay or misalignment, requiring calibration. The presented image-based processing could overcome those shortcomings of ECG gating. Additionally, our proposed method could be extended to compensate for other routine motions, such as respiratory motion gating, which conventional ECG is incapable of. While we use PA data for CG due to the inaccessibility to the concurrent US data, CG on US data may offer an advantage in its independence with the temporal variation of the PA contrast during ablation.

Despite the successful demonstration of the necrotic lesion visualization in the *in vivo* beating heart, we recognize several limitations to the current results and method. First, the ablation and its imaging were performed at the external surface of the heart instead of in an intracardiac environment. Our simplified experimental setup allowed us to focus on evaluating *in vivo* imaging capabilities by eliminating other factors, such as the difficulty of aligning the imaging setup with the ablation lesion, the inhomogeneous light illumination in the intracardiac space, and the light attenuation of a blood-filled condition. In addition, the shape and size of the ablation lesion should be controlled by the need to treat tissue in clinical practice instead of the linear geometry used in this study. Based on the outcome reported in the paper, we will focus on miniaturizing the imaging setup to validate the proposed imaging method in an intracardiac environment and to evaluate the imaging capability under the clinical workflow. Second, the current CG-sPA imaging pipeline does not grant a real-time visualization capability. Considering that a typical ablation process takes less than 1 minute [43–45], the current pipeline did not allow us to capture the tissue type transition in real-time. While the current *in vivo* CG-sPA system allows the intraoperative evaluation of the ablated tissue, real-time intra-ablation monitoring has not been realized. Increasing the imaging speed requires improving the SNR with less data duration. A high-speed laser with 100 Hz [46] or higher pulse repetition frequency increases the data points and can shorten the data recording time. Improving beamforming and the spectroscopic decomposition algorithm will also contribute to achieving real-time monitoring. Furthermore, the current spectroscopic decomposition was performed with reference spectra acquired from the *ex vivo* sample. A potential discrepancy between the *in vivo* and *ex* vivo tissue could result in a higher residual spectrum in our presented result. While our result has successfully detected the ablation-induced necrosis, using the spectra collected under the *in vivo* condition or including additional wavelengths for spectroscopic decomposition in the future work could improve the detection accuracy and access the oxygenation status better. Finally, the presented result was based on a single-lined ablation lesion with one swine sample, which limits the repeatability and statistical reliability. A study with a larger sample size is preferred and should be investigated as future work.

## 6. Conclusion

In this work, *in vivo* visualization of the cardiac ablation region at a beating heart was realized through CG-sPA imaging. Two light illumination modes were implemented to cross-validate the ablation lesion identification capability by using either three or sixteen wavelengths. Both modes successfully distinguished between the ablation-induced necrotic tissue and surrounding non-ablated tissue at the external surface of the heart. Hence, this *in vivo* demonstration supports the potential of PA imaging to be used for intraoperative cardiac ablation guidance.

## Disclosures

Tommaso Mansi was a scientist and employee of Siemens Healthineers USA during this work and currently is a scientist and employee of Janssen: Pharmaceutical Companies of Johnson & Johnson. Young-Ho Kim and Florin-Cristian Ghesu are scientists and employees of Siemens Healthineers USA. The other authors have no known competing financial interests or personal relationships that could have appeared to influence the work reported in this paper.

## Notes

### Competing Interest Statement

Tommaso Mansi is scientist and employee of Janssen: Pharmaceutical Companies of Johnson & Johnson. Young-Ho Kim and Florin-Cristian Ghesu are scientists and employees of Siemens Healthineers USA. The other authors have no known competing financial interests or personal relationships that could have appeared to influence the work reported in this paper.

### Summary of Updates

Additional analysis and discussion was included in the updated manuscript.

